# Early responses to hyperosmotic stress at the yeast vacuole

**DOI:** 10.1101/2025.08.11.669746

**Authors:** Kalaivani Saravanan, Patricia M. Kane

## Abstract

In yeast, early adaptation to hyperosmotic stress involves organelle-based mechanisms, including synthesis of phosphatidylinositol 3,5-bisphosphate (PI(3,5)P₂) at the vacuole. This low-level signaling lipid drives vacuolar fragmentation and activates the V-ATPase proton pump, which acidifies the vacuole and drives salt sequestration. The vacuole-resident V-ATPase subunit Vph1 interacts with PI(3,5)P₂ via its N-terminal domain (Vph1NT), directly linking lipid signaling to proton pump regulation. Under NaCl stress, PI(3,5)P₂ rapidly accumulates, triggering increased V-ATPase activity and vacuolar remodeling; these responses are impaired by deficient PI(3,5)P₂ synthesis. A Vph1NT-GFP fusion protein with no membrane domain is cytosolic without salt, but upon NaCl addition, rapidly relocalizes to a region adjacent to the vacuole in a PI(3,5)P2- dependent manner. The intensity and duration of this response depend on salt concentration. Vph1NT-GFP returns to the same location upon repeated salt challenge, suggesting that PI(3,5)P2 synthesis occurs at a localized domain/contact site. Disrupting PI(3,5)P₂ signaling, V-ATPase activity, or the high osmolarity glycerol pathway, which coordinates long-term transcriptional changes, compromises cellular adaptation to salt, underscoring the integration of lipid signaling and transcriptional regulation in hyperosmotic stress. These findings suggest activation of the V-ATPase, and possibly other targets, by PI(3,5)P2 synthesis provides immediate protection that primes cells for longer-term survival strategies.

**Significance Statement:** --Adaptation to high salt involves early responses at organelle membranes and slower transcriptional responses. The vacuolar/lysosomal signaling lipid, PI(3,5)P2 is critical for the early response, but the timing, localization, and targets of salt-induced PI(3,5)P2 synthesis are not fully understood.

--Experiments using Vph1NT-GFP as a low-affinity PI(3,5)P₂ biosensor suggest lipid synthesis occurs at a specific domain of the vacuolar membrane, with the level and duration of synthesis dependent on salt concentration and V-ATPase activity. A *hog1Δ* mutation ablates the slower response but elevates and extends PI(3,5)P2 activation.

--Controlled PI(3,5)P2 synthesis at the vacuole supports V-ATPase-driven salt sequestration; long-term adaptation requires both V-ATPases and the HOG pathway.

## INTRODUCTION

Cells must quickly adapt to changing extracellular environments that can induce multiple types of stress (Kültz, 2005; Hotamisligil and Davis, 2016). For example, hyperosmotic stress disrupts the balance of water and solutes across the plasma membrane, affecting intracellular ion balances, protein folding, organelle function, and cellular structure (Burg *et al*., 2007; Saito and Posas, 2012; Hohmann, 2015). NaCl stress requires cells to manage a toxic cation, as well as an osmotic imbalance (Ariño *et al*., 2010; Yang and Guo, 2018). Organisms have developed distinct but overlapping strategies to manage hyperosmotic stress. Osmoadaptation can be achieved by sensing extracellular solute concentrations and responding by counterbalancing the intracellular ion balance (Proft and Struhl, 2004), accumulating osmolytes (Blomberg and Adler, 1989), rearranging the cytoskeleton (Burg *et al*., 2007), sequestering salt into organelles (Xiong and Zhu, 2002; Ariño *et al*., 2010), and triggering transcriptional programs to achieve long-term osmoprotection (Hohmann, 2002; Kültz, 2005).

Plants, fungi, and certain animal cells are particularly vulnerable to hyperosmotic stress in their environments and have overlapping mechanisms of response (Hotamisligil and Davis, 2016). Hyperosmotic stress contributes to many human diseases, of which renal hyperosmolarity is a well-studied example (Yancey, 2005; Brocker *et al*., 2012). Altered osmolarity is associated with diabetes mellitus, dehydration, uremia, and hypernatremia (Burg *et al*., 2007; Brocker *et al*., 2012). In general, hyperosmotic shock triggers cell shrinkage, and adaptive mechanisms include the immediate activation of sensors that detect mechanical stress at the plasma membrane and cytoskeleton (Yancey, 2005; Burg *et al*., 2007; Christoph *et al*., 2007). Other early events include the activation of channels to regulate the flow of sodium and potassium ions and water, the sequestration of toxic cations in organelles such as the lysosome/vacuole, and the formation of stress granules (Proft and Struhl, 2004; Kültz, 2005; Hohmann, 2015). Ultimately, transcriptional activation supports increased synthesis of osmoprotectants and transporters to rebalance the stress and provide longer-term adaptation (Yancey, 2005; Christoph *et al*., 2007; Hohmann *et al*., 2007).

Plants and fungi subject to hyperosmotic stress, often accompanied by high salinity (salt stress), are protected by the accumulation of osmolytes and organic molecules such as proline, trehalose, and mannitol to mitigate water loss and maintain cellular integrity, along with modulation of water and ion transport and synthesis of stress-responsive proteins (Yancey, 2005; Hohmann *et al*., 2007). Plant and yeast vacuoles play critical roles in the response to acute osmotic stress (Gokbayrak *et al*., 2022). The yeast *S. cerevisiae* can tolerate high levels of hyperosmotic stress and can continue to grow in hyperosmotic media by employing both rapid response pathways and long-term adaptations to continued high salt conditions (Hohmann, 2002; Hohmann *et al*., 2007; de Nadal and Posas, 2022). Cell viability in high salt is compromised in mutants lacking organellar Na^+^/H^+^ exchangers (Nass and Rao, 1999; Cagnac *et al*., 2007), highlighting the importance of salt sequestration for cell survival. The high osmolarity glycerol (HOG) pathway, a stress-activated protein kinase cascade, directs metabolism toward osmolyte production at early times and later regulates gene expression, including channels that promote NaCl export, to restore cellular homeostasis (Hohmann, 2002; Saito and Posas, 2012). The transcriptional responses driven by the HOG pathway are slower but lead to long-term adaptation (Hohmann *et al*., 2007; Miermont *et al*., 2011). More immediately, yeast cells adjust the composition of their membrane lipids and sequester ions in the vacuole, the major site of salt storage, to help maintain cellular integrity (Herrera *et al*., 2013).

The phosphoinositide PI(3,5)P2 is a low-level signaling lipid that is enriched in late endosomes and lysosomes (Gary *et al*., 1998; Dove *et al*., 2002; Cullen and Carlton, 2012; Jin *et al*., 2016), including the yeast and plant lysosome-like vacuole. Levels of PI(3,5)P2 rise dramatically in response to NaCl-induced hyperosmotic stress (Dove *et al*., 1997; Duex *et al*., 2006b). The yeast Fab1 kinase converts PI(3)P to PI(3,5)P2 (Cooke *et al*., 1998; Gary *et al*., 1998) and functions in a complex with Vac14 (Bonangelino *et al*., 2002; Dove *et al*., 2002) and the PI(3,5)P2 phosphatase Fig4 (Rudge *et al*., 2004; Duex *et al*., 2006a). In yeast cells, the most obvious phenotype of PI(3,5)P2 depletion is very enlarged vacuoles (Bonangelino *et al*., 2002; Efe *et al*., 2007). Similarly, in mammalian cells, mutations result in cells with large vacuoles containing lysosomal markers LAMP1 and LAMP2 (Chow *et al*., 2007; Ferguson *et al*., 2009). In contrast, sustained high levels of PI(3,5)P2 synthesis in yeast result in highly fragmented vacuoles (Duex *et al*., 2006a; Lang *et al*., 2017), highlighting the role of this lipid in vacuolar morphology. Vacuolar fragmentation also occurs within minutes of acute salt stress and is dependent on PI(3,5)P2 (Zieger and Mayer, 2012).

V-ATPases are multi-subunit enzymes that use ATP hydrolysis to pump protons across organelle membranes, resulting in the acidification of vacuoles and lysosomes as well as other organelles (Collins and Forgac, 2020). The pH gradient generated by V-ATPases drives ions and nutrients across organelle membranes. Complete loss of V-ATPase activity is embryonically lethal in mammals (Sun-Wada *et al*., 2003) and conditionally lethal in yeast (Kane, 2007). V-ATPase structure is highly conserved across eukaryotes. V-ATPases consist of a membrane subcomplex, V0, that contains the proton pore, attached to a cytosolically oriented V1 subcomplex that contains the ATPase activity (Collins and Forgac, 2020). The largest membrane subunit, the V0 a-subunit, bridges the V1 and V0 subcomplexes via a large cytosolic N-terminal domain that is a major site of V-ATPase regulation (Tuli and Kane, 2023b). In yeast, the V0 a-subunit is present as two organelle-specific isoforms, Stv1 in the Golgi and Vph1 in the vacuole (Manolson *et al*., 1994). Loss of PI(3,5)P2 is linked to compromised vacuolar acidification (Gary *et al*., 1998; Bonangelino *et al*., 2002; Li *et al*., 2014).

Vacuolar vesicles isolated from *fab1Δ* mutants have approximately 50% of the ATPase activity of vesicles isolated from wild-type cells, along with a similar reduction in V1 subunit levels, suggesting that PI(3,5)P2 stabilizes V-ATPase assembly, as V0 subunit levels are maintained in the mutant vesicles (Li *et al*., 2014). Importantly, exposure of cells to 500 mM NaCl immediately before vacuole isolation almost doubles the V-ATPase activity in wild-type cells, and this increase is lost in vacuolar vesicles from salt-treated *fab1Δ* mutants (Li *et al*., 2014). The soluble N-terminal domain of yeast Vph1 (Vph1NT) binds to PI(3,5)P2 (Li *et al*., 2014; Banerjee *et al*., 2019), and the V0a1 isoform from humans also binds PI(3,5)P2 at a very similar sequence, suggesting that PI(3,5)P2 binding may be a general feature of V-ATPase regulation (Mitra *et al*., 2023).

We hypothesize that rapid PI(3,5)P2 synthesis activates Vph1-containing V-ATPases at the yeast vacuole, facilitating the uptake of salt into the vacuole through Na^+^/H^+^ exchangers and acting as part of the early response to salt stress before Hog1-dependent transcriptional responses are completed. In this study, we have used a GFP fusion to the cytosolic N-terminal domain of Vph1 (Vph1NT-GFP) in a microfluidic system to characterize the response to salt in real-time and analyze the dependence of the response on salt concentration, the localization of the initial response, and its relationship to the HOG pathway.

## RESULTS

### Vph1NT-GFP relocalization is dependent on salt concentration

Li et al. replaced the Vph1 C-terminal domain, which contains eight transmembrane domains, with a GFP tag (**Figure 1A**) and showed that the resulting Vph1NT-GFP fusion was recruited from the cytosol to a region adjacent to the vacuolar membrane after the addition of 0.5 M NaCl, conditions that transiently increase PI(3,5)P2 (Li *et al*., 2014). Membrane recruitment of Vph1NT-GFP was monitored through "snapshots” of different cell populations over time. However, with advances in microscopy and microfluidics techniques, we can now visualize responses in the same cells over time. Microfluidics allows us to continuously monitor cells maintained in a growth medium, before and after the addition of various salt concentrations, through time-lapse imaging. Our microfluidics setup is shown in **Figure 1A**. Cells suspended in medium without salt were immobilized on a concanavalin A-coated coverslip at the bottom of a chamber.

**Figure 1.**
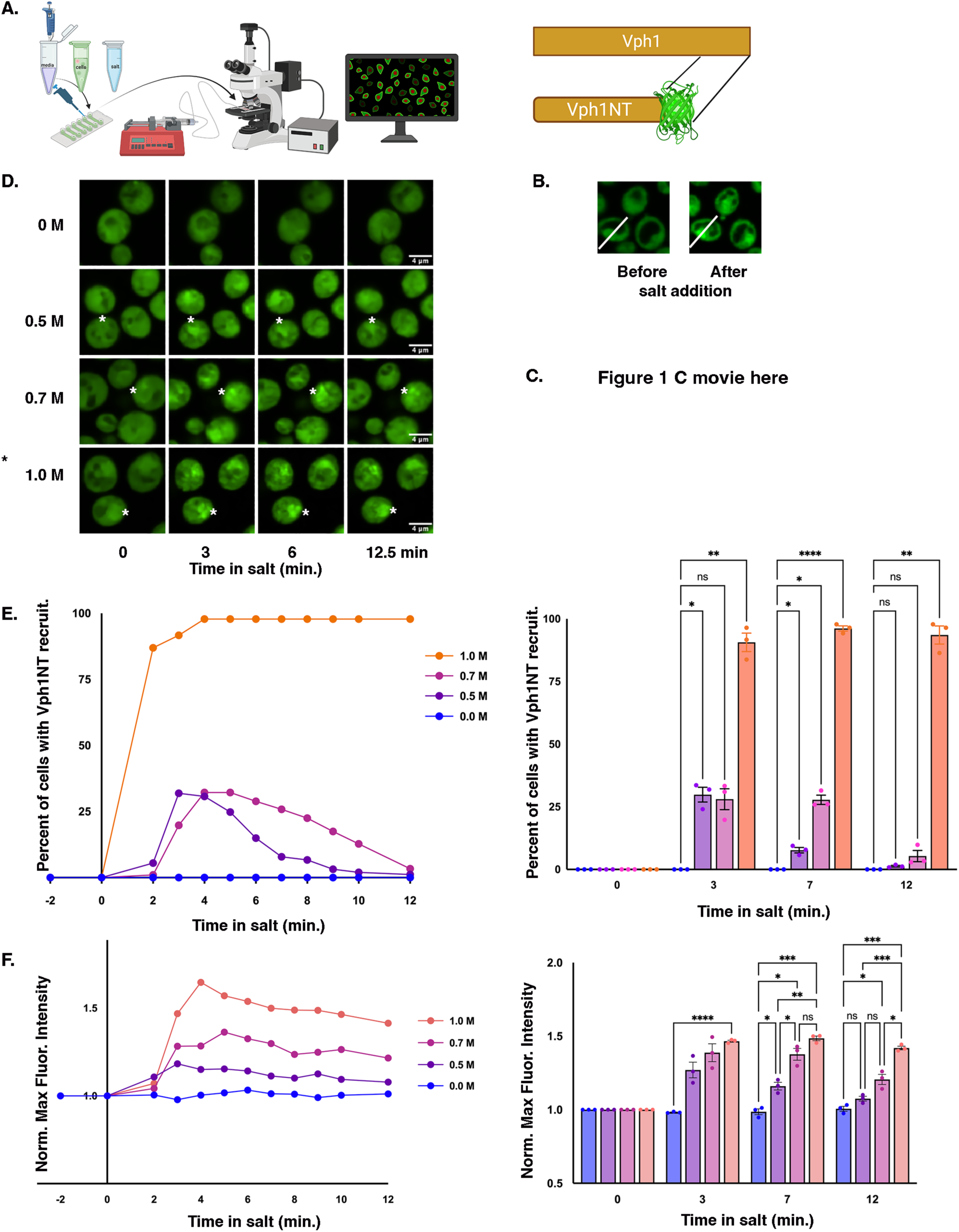
Vph1NT-GFP recruitment is dependent on salt concentration in haploid cells lacking V-ATPase activity. **(A) Left:** Schematic representation of the microfluidics setup used for live-cell imaging of yeast cells during salt treatment. **Right:** Schematic of the Vph1 full-length protein and the Vph1NT-GFP fusion, in which the C-terminal domain is replaced by GFP. **(B)** Representative example of line scans in the same cell before and after salt addition. Fluorescence intensity was measured along the yellow line. Scale bar = 4 μm. **(C)** Time-lapse video showing Vph1NT-GFP localization over time, before and after the addition of 0.7 M salt. All images were acquired as a single confocal section, and the images were taken every 30 seconds. **(D)** Time-lapse fluorescence microscopy of haploid yeast cells exposed to increasing salt concentrations. Time points shown are 0 min (no salt) and 3, 6, and 12.5 minutes after addition of the indicated salt concentrations. The white stars highlight Vph1NT-GFP relocalization near the vacuolar membrane in a single cell over time. Scale bar = 4 μm. **(E) Left:** Percentage of yeast cells exhibiting Vph1NT-GFP recruitment at different salt concentrations in a single biological replicate. For each salt concentration, the Vph1NT-GFP relocalization was assessed at each time in the same field of 250 to 300 cells and the percentage of total cells showing region(s) of enrichment of the Vph1NT-GFP signal was calculated. **Right:** Percentage of cells exhibiting Vph1NT-GFP recruitment at the indicated times and salt concentrations across three biological replicates. Each biological replicate (dot) represents an average from at least 250-300 cells, and error bars represent the mean ± SEM. ****P < 0.0001; ***P < 0.0005; **P < 0.005; *P < 0.05; ns = not significant were calculated by two-way ANOVA with repeated measures. **(F) Left:** Normalized maximum fluorescence intensity of Vph1NT-GFP obtained from line scans at different salt concentrations as described in Materials and Methods. Maximum fluorescence intensity was normalized to the average intensity before the salt addition at each point. Points shown are an average of 10-12 cells from a single biological replicate for each of the four salt concentrations. **Right:** Mean normalized maximum fluorescence intensity over time across three biological replicates. Each biological replicate (dot) represents results represents at least 10-12 cells, and error bars represent the mean ± SEM. ****P < 0.0001; ***P < 0.0005; **P < 0.005; *P < 0.05; ns = not significant were calculated by two-way ANOVA with repeated measures.

Li et al. (2014) demonstrated Vph1NT-GFP recruitment in haploid yeast cells only at 500 mM salt. We sought to repeat those measurements and assess the effect of varied salt concentrations in the microfluidic system in real time. GFP fluorescence in the immobilized cells was initially monitored for at least three minutes. Time-lapse imaging was continued for control samples, but for salt-treated samples, medium containing the indicated concentration of NaCl was then pumped into the chamber, and the same field was imaged over time. An area of increased fluorescence intensity adjacent to the vacuole appeared shortly after salt addition in salt-treated cells but was not seen without salt treatment, as shown in **Figure 1B** and a time-lapse video of cells treated with 700 mM salt **(Video 1C).** Yeast cells from select time points after the addition of the indicated salt concentrations are compared in **Figure 1D**. We analyzed the Vph1NT-GFP response over time in two ways. First, we counted the number of cells with visible recruitment over time at each salt condition. The percentage of cells showing Vph1NT-GFP recruitment varied with salt concentration, with less than 50% of cells showing any accumulation after treatment with 500 mM and 700 mM salt, while almost all cells responded to 1 M salt exposure (**Figure 1E**). This suggests there is a threshold for the response that is dependent on salt concentration. Second, we monitored the intensity and duration of GFP relocalization by line scan through the region of accumulated GFP fluorescence in the same cells over time (see Figure 1B), quantified the maximal fluorescence at each time point, and then normalized to the fluorescence before salt addition. **Figure 1F** shows the normalized maximal fluorescence intensity over time for a single representative experiment, as well as the mean responses at the indicated times across three biological replicates. These experiments confirm the relocalization of Vph1NT-GFP in response to salt, observed by (Li *et al*., 2014), and extend that observation to demonstrate that the proportion of cells that respond, the intensity of the maximal fluorescent signal, and the duration of the response are all salt concentration-dependent.

### Vph1NT-GFP relocalization in cells with active vacuolar ATPase

Haploid yeast cells containing Vph1NT have no V-ATPase activity at the vacuole because, in the absence of a membrane-bound Vph1, V0 subcomplexes containing Vph1 do not assemble (Hill and Stevens, 1995; Kawasaki-Nishi *et al*., 2001). The smaller population of Stv1-containing V0 complexes generally does not reach the vacuole, and Stv1NT does not bind PI(3,5)P2 (Manolson *et al*., 1994; Tuli and Kane, 2023a). Although the V1 subcomplex is assembled and cytosolic, its ATPase activity is silenced, and it does not bind to Vph1NT (Parra *et al*., 2000; Oot *et al*., 2017). Lack of V-ATPase activity strongly affects the cellular response to salt stress (Li *et al*., 2012), and we reasoned that it could affect Vph1NT-GFP recruitment in response to salt. To address this possibility, we generated a heterozygous diploid strain containing one copy of wild-type *VPH1* and one copy of Vph1NT-GFP. These cells should have 50% of the wild-type level of V-ATPase at the vacuole, a level sufficient to suppress most growth defects arising from the loss of vacuolar V-ATPase activity (Diakov and Kane, 2010). We exposed the diploid cells to different salt concentrations in the microfluidic system, as described in Figure 1. Images of cells at different time points with varied salt concentrations are shown in **Figure 2A**. A time-lapse video of Vph1NT-GFP before and after treatment with 700 mM salt is shown in **Video 2B**. As in the haploid strain, Vph1NT-GFP is cytosolic in the heterozygous diploid cells in the absence of salt (**Figure 2A**). The percentage of cells showing Vph1NT-GFP relocalization was again salt concentration-dependent, but the percentage of cells responding to 500 mM and 700 mM salt was much higher than in haploid cells (**Figure 2C**). At 1 M salt, recruitment was again observed in 100% of cells. Figure 2C shows a representative time course from a single experiment (left) as well as the average of three biological replicates at the indicated times (right). The changes in fluorescence intensity with time followed the same trend as in haploid cells. However, the fluorescence accumulation after salt stress appears to be more intense in **Figure 2A**, and this is supported by the normalized fluorescence intensity measurements in **Figure 2D**, where the normalized peak fluorescence intensity almost doubled in cells treated with 700 mM salt. These data suggest that there is a lower threshold and a more robust Vph1NT-GFP response in cells that retain V-ATPase activity in the vacuole.

**Figure 2.**
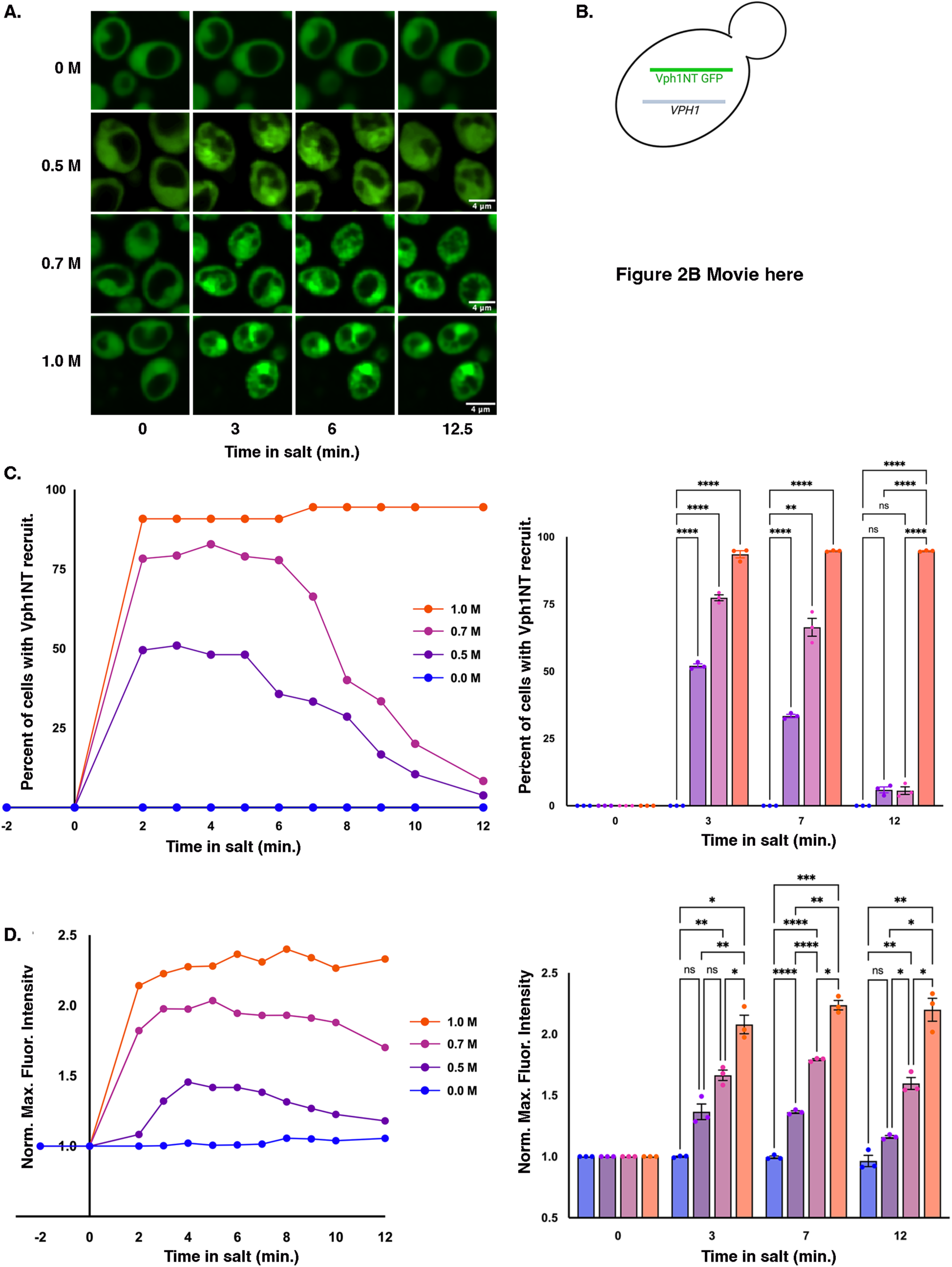
Salt-dependent recruitment of Vph1NT-GFP in the presence of active V-ATPases. **(A)**Time-lapse images of heterozygous diploid yeast cells (*VPH1*/Vph1NT-GFP) before and after the addition of the indicated salt concentrations. Relocalization of Vph1NT-GFP is shown at 0 min, 3 min, 6 min, and 12.5 min. Scale bar = 4 μm. **(B)** Schematic of a diploid yeast cell heterozygous for full-length Vph1 (Vph1FL) and Vph1NT-GFP. **(C)** Time-lapse video showing Vph1NT-GFP localization over time, before and after the addition of 0.7 M salt. All images were acquired as a single confocal section, and images shown were taken at 30-second intervals. **(D)** The percentage of yeast cells exhibiting Vph1NT-GFP recruitment at different salt concentrations in the presence of active V-ATPase was determined as in Figure 1E. Left: The Percentage of yeast cells exhibiting Vph1NT-GFP recruitment at different salt concentrations in a single experiment (biological replicate) was determined as described in Figure 1E. Right: Percentage of cells with Vph1NT-GFP relocalization at the indicated times and salt concentrations across three biological replicates as determined in Figure 1E. Each biological replicate (dot) represents at least 250-300 cells, and error bars represent the mean ± SEM. ****P < 0.0001; ***P < 0.0005; **P < 0.005; *P < 0.05; ns = not significant. Significance was calculated by two-way ANOVA with repeated measures. **(E)** Normalized maximum fluorescence intensity of Vph1NT-GFP at different salt concentrations in the presence of active V-ATPase was determined as described in Figure 1F. Left: Normalized maximum fluorescence intensity for a single biological replicate of 10-12 cells. Right: Average normalized maximum fluorescence intensity at the indicated times across three biological replicates. Each biological replicate (dot) represents at least 10-12 cells, and error bars represent the mean ± SEM. ****P < 0.0001; ***P < 0.0005; **P < 0.005; *P < 0.05; ns = not significant. Significance was calculated by two-way ANOVA with repeated measures.

### Vph1NT-GFP relocalization in response to salt stress is PI(3,5)P2-dependent

We confirmed that Vph1NT recruitment is PI(3,5)P2-dependent by two different approaches. We generated a homozygous *vac14Δ/vac14Δ* mutation in the heterozygous diploid containing one copy each of *VPH1* and Vph1NT-GFP. *vac14Δ* mutants have much lower levels of PI(3,5)P2 than wild-type cells and, like other mutants lacking PI(3,5)P2, have greatly enlarged vacuoles (Zhang *et al*., 2007; Jin *et al*., 2008). We saw some heterogeneity in vacuole size, as described previously (Rudge *et al*., 2004; Jin *et al*., 2008), and rapid suppression of the *fab1Δ* phenotype has been reported (Jin *et al*., 2017). To avoid any suppressors, we selected cells with enlarged vacuoles and quantified the Vph1NT-GFP response to 700 mM salt (**Figure 3A**). Quantitation of normalized maximal fluorescence intensity in **Figure 3B** confirms significantly less relocalization of Vph1NT-GFP in the *vac14Δ* cells vs. wild type. The small increase in fluorescence may be attributed to the shrinkage of the cells in the salt (Hohmann, 2002) or the small amount of PI(3,5)P2 present in *vac14* mutants (Bonangelino *et al*., 2002). These data confirm that PI(3,5)P2 is required for Vph1NT-GFP relocalization in response to NaCl stress.

**Figure 3.**
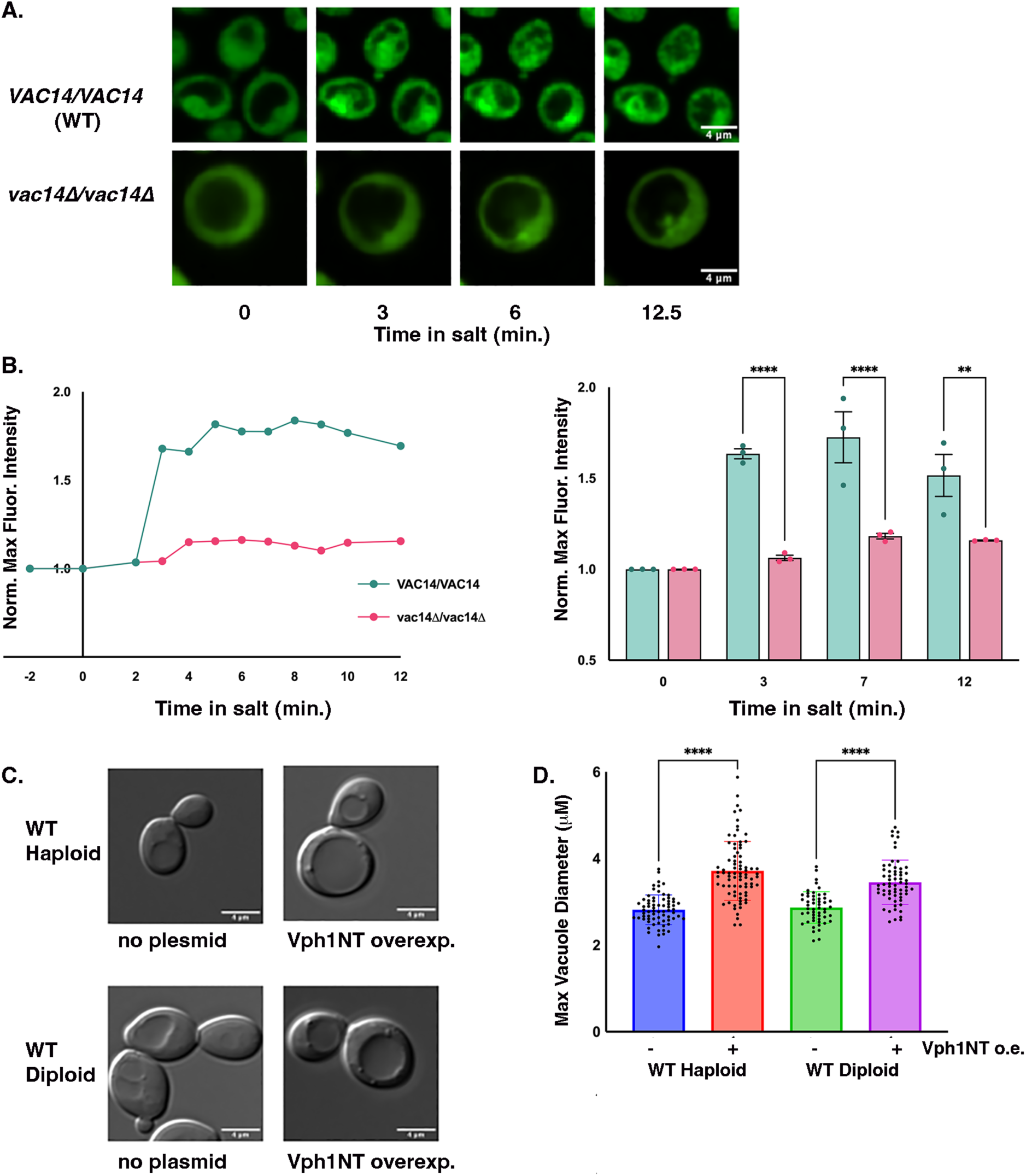
Vph1NT-GFP re-localization in response to salt stress is dependent on PI(3,5)P₂. **(A) Top:** *VAC14/VAC14* (WT) diploid cells exposed to 0.7 M salt for 0-, 3-, 6-, and 12.5 min. **Bottom:** Homozygous *vac14Δ/vac14Δ* diploid cells subjected to the 0.7 M salt treatment for the same time periods. Scale bars: 4 µm. **(B) Left**: Normalized maximum fluorescence intensity of Vph1NT-GFP in *VAC14/VAC14* and *vac14Δ/vac14Δ* cells across the indicated times. **Right:** Quantification of the maximum fluorescence intensity across three biological replicates for WT and *vac14Δ/vac14Δ* cells performed as described in Figure 1F and Materials and Methods. Each dot represents a different biological replicate of at least 10-12 cells, and error bars represent the mean ± SEM. ****P < 0.0001; **P < 0.005; calculated by multiple t-test. **(C)** Differential interference contrast (DIC) images of WT haploid and diploid cells with and without Vph1NT overexpression illustrate changes in vacuole morphology. Scale bars: 4 µm. **(D)** The largest vacuole diameter in haploid WT and diploid WT cells, with and without Vph1NT overexpression, was measured in 50-75 cells per biological replicate. Dots represent all vacuolar diameters quantitated across the three replicates along with the mean ± SEM. ****P < 0.0001, calculated by unpaired Student’s t-test.

We would also predict that if Vph1NT-GFP is binding directly to PI(3,5)P2, then Vph1NT overproduction might saturate the very limited pool of PI(3,5)P2, resulting in PI(3,5)P2 deficiency phenotypes such as vacuole enlargement. Recent data suggest that several widely used PIP sensors compete with native PIP binding proteins and generate phenotypes suggestive of PIP depletion (Holmes *et al*., 2025). To test this possibility, we expressed Vph1NT from a multicopy plasmid in haploid and diploid wild-type cells. Vacuoles, visualized as indentations under DIC optics, appear larger in both haploid and diploid cells overexpressing Vph1NT from the plasmid (**Figure 3C**). Vacuole size was quantified by measuring the diameter of the largest vacuole in 50- 70 different haploid or diploid cells in each of the 3 biological replicates. The results are quantified in **Figure 3D** and indicate a significant increase in vacuolar diameter when Vph1NT is overexpressed. These data suggest that Vph1NT overexpression can generate PI(3,5)P2 deficiency phenotypes by competing with endogenous proteins for lipid binding (Holmes *et al*., 2025). Taken together, these data are consistent with direct binding of Vph1NT to PI(3,5)P2 and indicate that the relocalization of Vph1NT-GFP in response to salt stress is PI(3,5)P2-dependent.

We tested whether genomically integrated Vph1NT-GFP, expressed from the *VPH1* promoter (Figure 1), alters vacuole size under basal conditions, where PI(3,5)P2 levels are very low. As shown in **Supplemental Figure 1**, the mean vacuole diameter of the haploid cells expressing Vph1NT-GFP (without salt) was the same as the diameter of a haploid *vph1Δ* mutant (which also lacks an intact V-ATPase). Thus, Vph1NT does not significantly deplete available PI(3,5)P2 when present in a single copy.

### Temporal and spatial responses to repeated salt stress

Induction of PI(3,5)P2 synthesis in response to salt is a rapid, transient response (Dove *et al*., 1997). PI(3,5)P2 levels were previously shown to increase 20-fold upon the addition of 0.9 M NaCl and return to baseline within 20 minutes (Duex *et al*., 2006a), consistent with the timing of Vph1NT-GFP relocalization shown in Figures 1 and 2. Given this dynamic response, we asked whether Vph1NT-GFP responds similarly when salt is added a second time. To address this, we again used the heterozygous (*VPH1*/Vph1NT-GFP) diploid cell. **Figure 4A** shows the strategy and timing for the first and the second salt additions. After initial imaging with no salt, we added 700 mM salt and continued monitoring until the response had largely reversed (13 minutes after salt addition). We then added medium without salt for an additional 7 min. During this time, Vph1- GFP fully returned to the cytosol. We repeated the addition of 700 mM NaCl and continued to image Vph1-GFP as shown in **Figure 4B**. Vph1NT-GFP relocalized with similar kinetics after the second addition of salt. The peak fluorescence intensity did not appear to be as high as after the first addition, but the peak intensities after the first and second salt additions were not significantly different when analyzed across replicates (**Figure 4C**). To control for bleaching of Vph1-GFP over the extended time of the experiment, we also monitored Vph1-GFP fluorescence over the same total time in cells never treated with salt and observed no decrease in signal. The results suggest that the cells respond to a second salt addition, even when it is applied after a relatively short time. There may be some desensitization of the response, but if so, the mechanistic basis will require further exploration.

**Figure 4.**
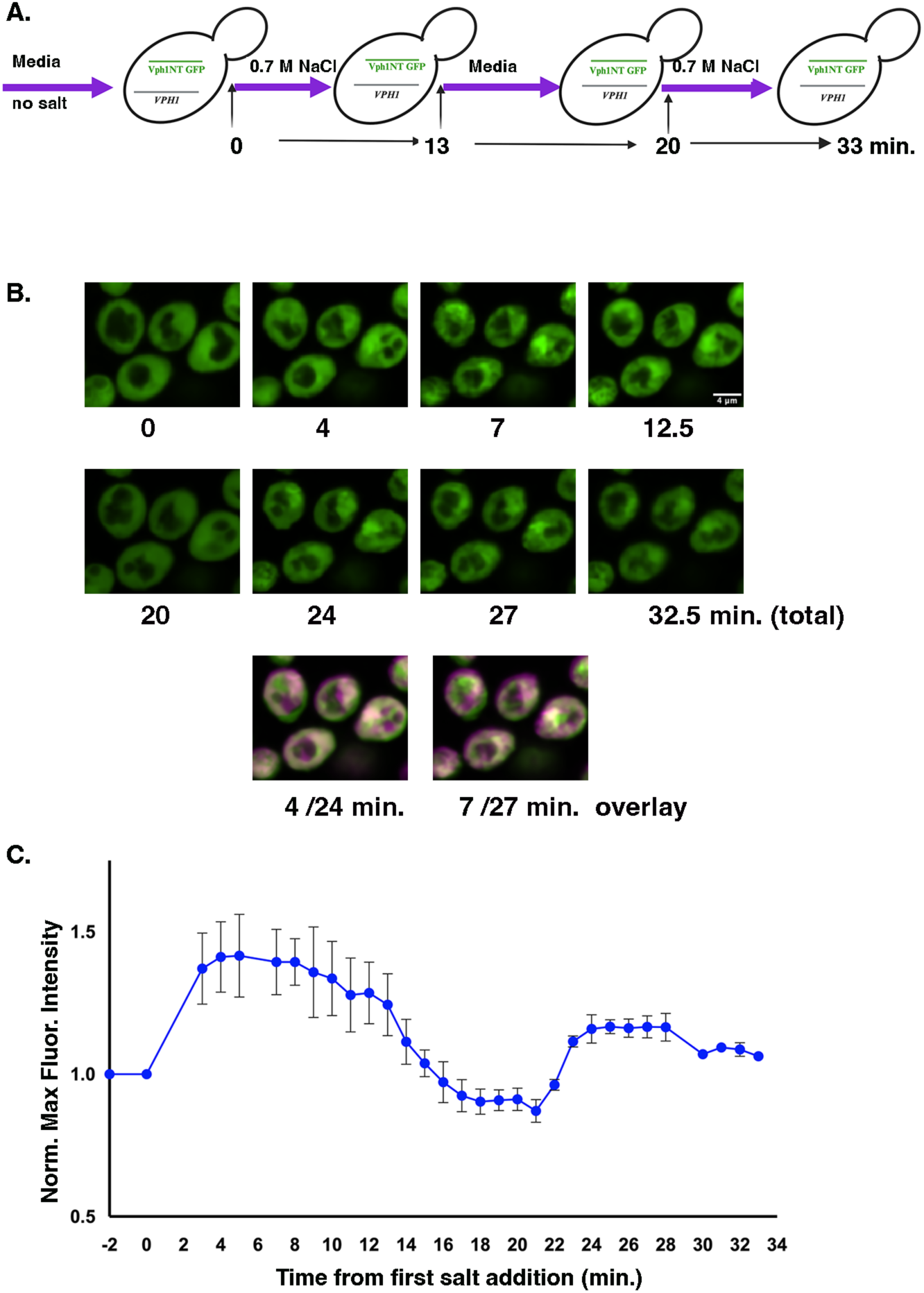
Vph1NT-GFP relocalization upon repeated salt exposure. **(A)**Schematic of the experimental timeline, indicating the timing of media addition, salt addition, media re-addition, and salt re-addition. (**B)** Images of Vph1NT-GFP from VPH1/Vph1NT-GFP diploid cells. Top: Images captured during the initial exposure to 0.7 M salt. Representative time points (0, 4, 7, and 12.5 minutes) are shown. Middle: Images of the same cells following a 7- minute recovery period, followed by a second addition of 0.7 M salt. The second salt addition occurred at the 20 min. time point. Bottom: images from 4 min and 7 min after the first salt addition (green) and second salt addition (magenta) are overlaid to highlight the return of Vph1NT to the same position. **(C)** Normalized maximum fluorescence intensity of Vph1NT-GFP over the entire time course of the experiment (diagrammed in Figure 4A). The mean + the range of two biological replicates is plotted. Maximum fluorescence intensities throughout the time course were normalized to the intensity before the first salt addition. Each biological replicate represents at least 10-12 cells.

We see a single focused area of Vph1NT-GFP fluorescence after salt addition in most cells (**Figures 1 and 2**). We were surprised to find that Vph1NT-GFP returns to the same location after repeated salt addition, as indicated by the overlay of images 4 and 7 min after the first and second salt additions (**Figure 4B**, **bottom**). This suggests that the location of the recruitment is not random but occurs at a specific area of the vacuole or membrane domain. One advantage of using heterozygous diploid cells is that we can follow both Vph1NT-GFP and full-length Vph1 labeled with mCherry (Vph1-mCherry) at the same time. Images of Vph1NT-GFP/Vph1-mCherry heterozygous diploid cells before and after the addition of 700 mM NaCl are shown in **Figure 5A** and the video in Figure **5B**. Vph1-mCherry outlines the vacuolar membrane, but also clearly shows invagination of the vacuolar membrane after salt addition, as it progresses toward vacuole fragmentation. As described above, Vph1NT-GFP does not recruit to the entire vacuolar membrane upon salt addition; in most cells, it recruits to the site of membrane invagination after salt addition. In some images, this area appears to be partially depleted of Vph1-mCherry. The merged images and the time-lapse video in **Figures 5A** and **B** show some limited overlap of Vph1-mCherry and Vph1NT-GFP in and immediately around this region at early times after salt addition, which is dispersed after longer times as Vph1NT-GFP returns to the cytosol.

**Figure 5.**
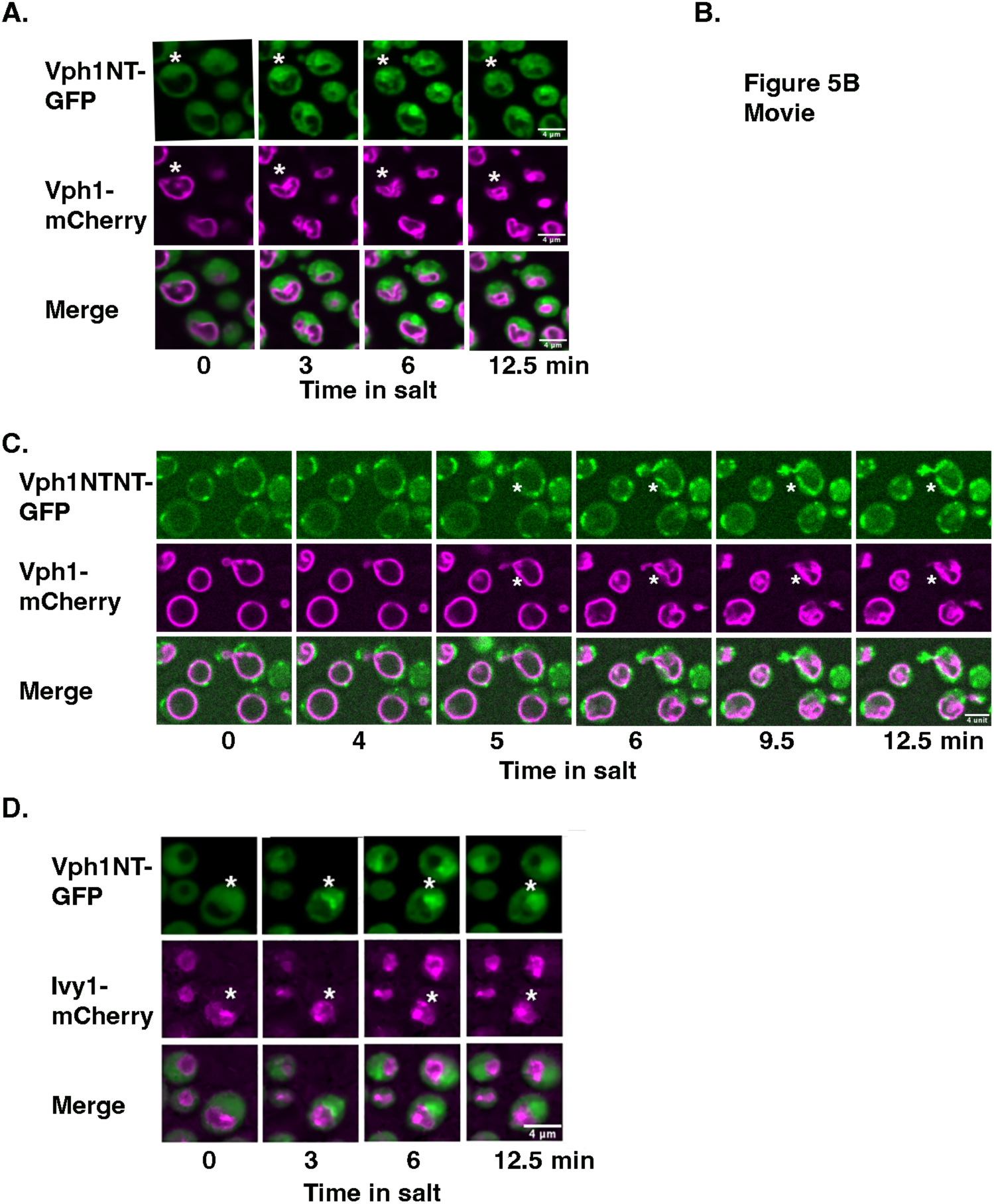
Both Vph1NT and the duplicated Vph1NTNT localize to regions of vacuolar invagination after salt addition. **(A)** Time-lapse fluorescence microscopy of diploid yeast cells heterozygous for Vph1NT-GFP (green, top panel) and full-length Vph1-mCherry (magenta, middle panel) exposed to 0.7 M NaCl. Images shown were obtained 0, 3, 6, and 12.5 minutes after salt addition. The bottom panel shows merged images of GFP and mCherry signals. The white star highlights a region of the vacuole undergoing invagination after salt addition, demonstrating the dynamic recruitment of Vph1NT-GFP to this region. Scale bar = 4 μm. **(B).** A time-lapse video of the merged images over the entire time course. Images were from a single confocal plane and were acquired every 30 seconds. **(C) Top panel:** Images of diploid cells expressing Vph1NTNT-GFP at 0, 4, 5, 6, 9.5, and 12.5 minutes after exposure to 0.7 M NaCl. **Middle panel:** Corresponding images of the same cells expressing Vph1FL-mCherry (magenta). **Bottom panel:** Merged channels. Scale bar = 4 μm. White stars indicate a vacuole undergoing invagination after salt addition. **(D)** Images of diploid cells heterozygous for both Vph1NT-GFP and *VPH1* and Ivy1-mCherry and *IVY1*. **Top panel:** Vph1-GFP (green) at 0-, 3-, 6-, and 12.5-min addition of 0.7 M NaCl. **Middle panel:** Images of Ivy1-mCherry in the same cells (magenta). **Bottom panel:** Merged channels. Scale bar = 4 μm.

Huda et al. recently visualized the response of a yeast SnxA-GFP PI(3,5)P2 sensor to the addition of 900 mM NaCl (Huda *et al*., 2024). They also observed transient relocalization of SnxA-GFP to a focused region of the vacuolar membrane with a time course similar to that seen for Vph1NT-GFP in Figure 1. They frequently saw that SnxA-GFP was recruited to sites of vacuolar invagination/fission. SnxA is likely to have a higher affinity for PI(3,5)P2 than Vph1NT (Vines *et al*., 2023). We previously generated a higher avidity probe by duplicating Vph1NT (Vph1NTNT- GFP) and reported that it showed both some baseline recruitment to the vacuolar membrane and a longer duration of vacuolar localization in response to 500 mM NaCl (Li *et al*., 2014). We introduced a copy of Vph1NTNT-GFP into the diploid strain, generating a Vph1- mCherry/Vph1NTNT-GFP heterozygote, and visualized the response to the addition of 700 mM NaCl in the microfluidics system. As shown in **Figure 5C**, Vph1NTNT-GFP is localized to both small patches and the cytosol in the absence of salt. We begin to see the brightening of Vph1NTNT-GFP patches on the vacuolar membrane, as well as invagination of the vacuolar membrane within 5 min. of salt addition. As indicated in Figure 5C (stars), there are patches of Vph1NTNT-GFP at sites that are folding in toward the vacuolar lumen. In addition, consistent with a higher affinity of the Vph1NTNT-GFP probe as a result of Vph1NT duplication, vacuolar localization persists longer, in some cases resulting in the probe encircling the vacuolar membrane at times when the Vph1NT-GFP signal at the vacuole has declined (12.5 min). These results suggest that the relatively low affinity of Vph1NT-GFP supports the detection of the peak of PI(3,5)P2 synthesis in the first few minutes after salt addition, but as the lipid diffuses across the vacuolar membrane, its concentration becomes too low to sustain Vph1NT-GFP recruitment. In contrast, vacuolar localization of Vph1NTNT-GFP remains as PI(3,5)P2 diffuses away from the site of initial synthesis.

Fab1 contains the catalytic site required for PI(3,5)P2 synthesis and is localized around the vacuole (Gary *et al*., 1998). However, there is evidence that Fab1 may be inhibited by interaction with Ivy1 and that colocalization of Fab1 and Ivy1 is reduced in the presence of salt stress, allowing activation of PI(3,5)P2 synthesis (Malia *et al*., 2018). We hypothesized that salt-induced PI(3,5)P2 synthesis and Vph1NT-GFP recruitment might occur at the vacuolar regions cleared of Ivy1. We visualized the localization of Ivy1-mCherry and Vph1NT-GFP before and after the addition of 700 mM salt (**Figure 5D**). Ivy1 generally localizes to the vacuolar membrane in the absence of salt, as reported previously (Malia *et al*., 2018). However, by 3 min after the salt addition, Vph1NT-GFP and Ivy1-mCherry are segregated into two different regions of the vacuole. These results are also consistent with Vph1NT-GFP marking sites of PI(3,5)P2 synthesis, where Ivyq has been removed, in salt-treated cells.

### The duration of Vph1NT-GFP recruitment is increased in a *hog1Δ* mutant

Under salt stress, the proton gradient established by the V-ATPase drives the uptake of Na^+^ into the vacuole through Na^+^(K+)/H^+^ exchangers (Nass and Rao, 1999; Cagnac *et al*., 2007; Li *et al*., 2012). This is a rapid response to salt stress that we hypothesize is enhanced by increased levels of PI(3,5)P2. In contrast, the HOG pathway provides a sustained transcriptional response to stress that parallels vacuolar accumulation, but full transcriptional activation is slower (Pelet and Peter, 2011; Jin *et al*., 2017). Deletion of *VMA* genes and *HOG1* leads to synthetic growth defects, even at relatively low salt concentrations, and the *vma* mutants even show some activation of the HOG pathway in the absence of salt (Li *et al*., 2012). We integrated Vph1NT- GFP into a haploid *hog1Δ* mutant and imaged the cells before and after the addition of 700 mM NaCl as described above. Preliminary experiments suggested that Vph1NT-GFP might relocalize for longer times in the *hog1Δ* mutant, so we conducted imaging for 27 min. after salt addition (**Figure 6A**). As shown in **Figure 6B**, the normalized maximum intensity of Vph1NT-GFP was higher in the *hog1Δ* mutant by 5 min after salt addition, and even more strikingly, was sustained above the initial level through 25 min, well after the intensity of Vph1NT-GFP had returned to baseline in wild-type cells. These results suggest that levels of PI(3,5)P2 synthesis are higher and are sustained for longer times in the absence of Hog1 activity. Very similar results were obtained in a diploid strain that was homozygous for the *hog1Δ* mutant, but heterozygous for Vph1NT- GFP/*VPH1*. The higher, sustained level of PI(3,5)P2 would be expected to activate the V-ATPase (Li *et al*., 2014) and might help to compensate for the loss of Hog1, particularly in the diploid cell, which contains intact V-ATPases.

**Figure 6.**
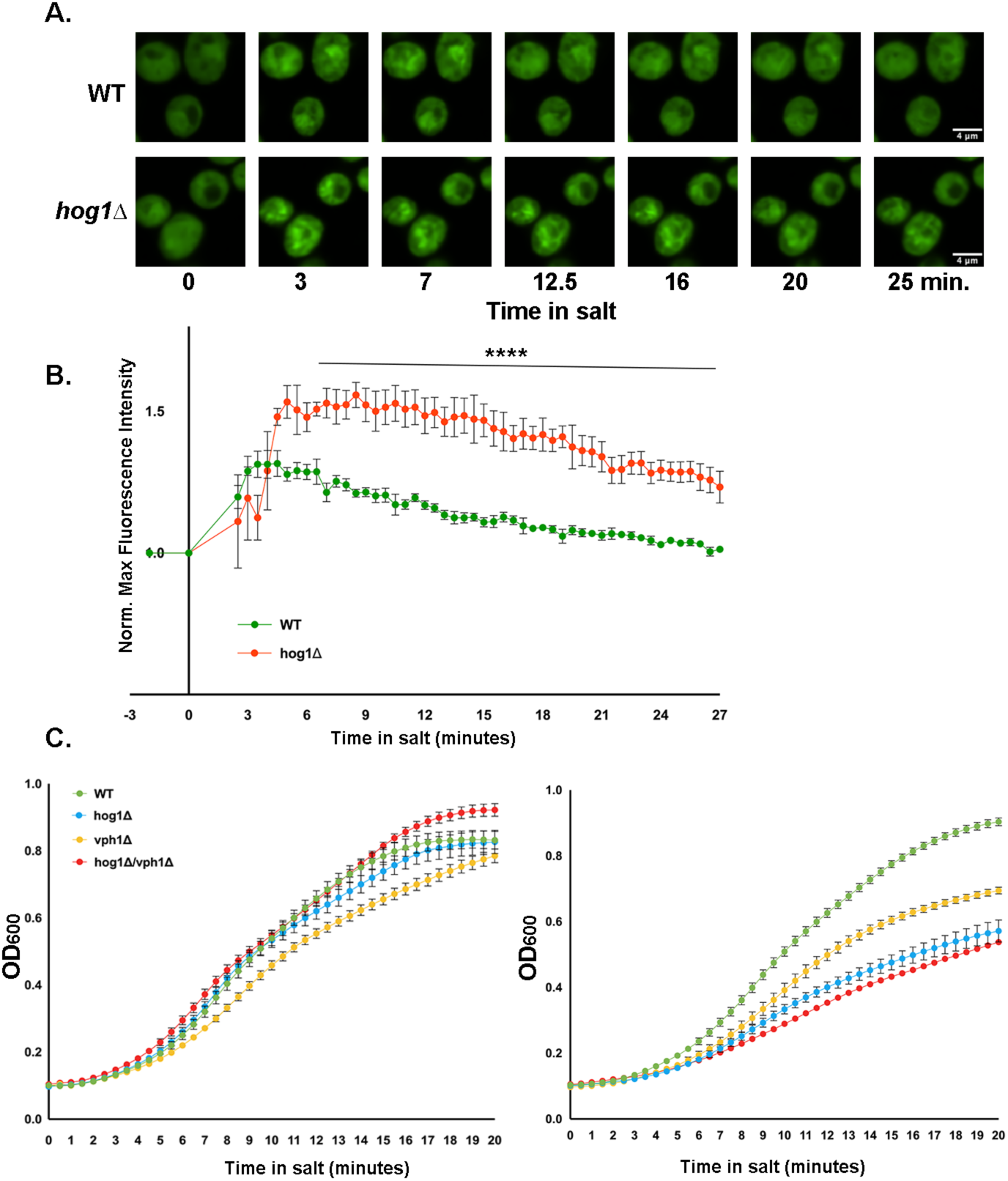
Vph1NT-GFP localization at the vacuolar membrane is prolonged in the absence of Hog1, but both Vph1 and Hog1 are required for long-term salt adaptation. **(A)** Fluorescence microscopy of haploid WT (top panel) and *hog1Δ* (bottom panel) cells expressing Vph1NT-GFP following treatment with 0.7 M NaCl. Time points shown include 0, 3, 7, 12.5, 16, 20, and 25 minutes. **(B)** Normalized maximum fluorescence intensity of Vph1NT-GFP was determined at the indicated time points in haploid WT and *hog1Δ* cells under 0.7 M NaCl treatment as described in Figure 1F. Data from three biological replicates (10-12 cells each) were normalized to the pre-salt treatment intensity for each replicate. Mean ± SEM is shown. A t-test with multiple comparisons determined a P-value <0.0001 (****) for the indicated time range. **(C)** Growth curves for WT, *hog1Δ*, *vph1Δ*, and *hog1Δvph1Δ* double mutant in the absence of salt (left) and the presence of 500 mM NaCl (right) are shown. Mean ± SEM for three biological replicates is shown for each curve.

Uptake of Na^+^ ions through vacuolar antiporters, supported by salt-induced increases in PI(3,5)P2 that activate the V-ATPase and maintain the pH gradient, is likely to play a critical short-term role in the survival of cells exposed to hyperosmotic stress. However, the vacuole does not have an unlimited capacity for salt sequestration, so longer-term mechanisms, such as transcriptional activation of Na^+^ export channels via the HOG MAPK pathway, are critical for cells to survive prolonged hyperosmotic stress (Ariño *et al*., 2010). To address the relative contributions of V-ATPase activity at the vacuole and the HOG pathway over longer times, we compared the growth of wild-type cells and *vph1Δ, hog1Δ, and vph1Δhog1Δ* mutants over 20 hours at 0- and 500-mM salt. As shown in **Figure 6C**, the wild-type and mutant strains have similar growth rates in the absence of salt. Growth of the *vph1Δ* mutant is slowed in the presence of 500 mM NaCl, consistent with an important role in vacuolar V-ATPase activity in the presence of salt stress. However, the growth rate of the *hog1Δ* mutant is affected more strongly, and there is no significant difference between the growth of *hog1Δ* and *hog1Δvph1Δ* mutants after 20 hours. These data highlight the importance of Hog1 activity over prolonged salt stress. Extending the accumulation of PI(3,5)P2 at the vacuole in the *hog1Δ* mutant, as seen in Figures 6A and B, may provide a short-term compensation but does not appear to be sufficient for long-term osmoadaptation.

## DISCUSSION

PIP-binding specificity is characteristic of several of the V-ATPase a-subunit isoforms from both yeast and human cells (Li *et al*., 2014; Banerjee *et al*., 2019; Mitra *et al*., 2023; Tuli and Kane., 2023b). The *in vitro* data from liposome pelleting assays suggest that purified Vph1NT binds relatively weakly to PI(3,5)P2 relative to the binding of the other V0-aNT domains to their cognate PIP lipids (Tuli and Kane, 2023a, 2023b). However, like the aNT domains of the several other a-subunit isoforms (Banerjee *et al*., 2019; Mitra *et al*., 2023), expressed Vph1NT-GFP relocalizes from the cytosol to membranes containing specific PIP lipids, in this case, PI(3,5)P2. We build on these results here and propose that Vph1NT-GFP is a low-affinity sensor for PI(3,5)P2, particularly suitable for detecting dynamic changes in PI(3,5)P2 levels at the vacuole at early stages of salt stress in real-time. By monitoring Vph1NT-GFP in a microfluidic system, we demonstrate that the duration and intensity of Vph1NT-GFP relocalization are dependent on salt concentration and that the intensity and threshold of the response are both increased in cells with active V-ATPases at the vacuole. Consistent with previous evidence that PI(3,5)P2 levels are increased transiently (Dove *et al*.,1997; Duex *et al*., 2006a), Vph1NT-GFP returns to the cytosol even when extracellular salt concentrations remain high. However, after the removal of extracellular salt for only a few minutes followed by readdition, Vph1NT-GFP again moves from cytosol to the same region peripheral to the vacuole, a location where invagination of the vacuolar membrane occurs early in the process of vacuolar fragmentation. These data provide novel insights into the early, PI(3,5)P2-dependent events at the vacuole that occur in response to salt stress.

The kinetics of Vph1NT-GFP relocalization after the addition of 700 mM and 1.0 M NaCl (Figures 1 and 2) are consistent with the kinetics of the rise and fall of PI(3,5)P2 levels measured in response to 900 M NaCl (Duex *et al*., 2006a). However, the salt concentration dependence of PI(3,5)P2 levels had not been explored in depth. In Figures 1 and 2, the initial response to 500 mM NaCl is still rapid but is both shorter-lived and observed in a small proportion of the cell population. Interestingly, the initial phosphorylation of Hog1 in response to salt addition, which redirects metabolism toward glycerol synthesis, shows a graded response to changes in concentration, while the transcriptional response exhibits a “biphasic” concentration-dependent threshold in which not all cells respond (Pelet and Peter, 2011). The more robust response in diploid cells that have V-ATPase activity at the vacuole, compared to haploids that do not, was somewhat unexpected. We previously showed that PI(3,5)P2 synthesis in response to salt stress activates V-ATPase activity, so the diploid cell should have an increased capacity to sequester Na^+^ in the vacuole relative to the haploid, with no V-ATPase activity at the vacuole. Theoretically, this could decrease the level of salt stress, but the results suggest that PI(3,5)P2 synthesis may be increased in the presence of V-ATPase activity and/or vacuolar acidification. Alternatively, compensatory mechanisms that are activated in the absence of V-ATPase activity at the vacuole may blunt the PI(3,5)P2 response (Li *et al*., 2012).

We find that Vph1NT-GFP accumulates in a specific location adjacent to the vacuolar membrane, both upon initial exposure to high salt and on a subsequent shift to salt (Figures 4B and 5A). In some images, this location appears to be partially depleted of full-length Vph1 even before salt addition (Figure 5A), and after addition, the membrane begins to invaginate into the lumen at this site. Other PI(3,5)P2 sensors also exhibit asymmetric recruitment to the vacuolar membrane upon salt stress. Huda et al. (Huda *et al*., 2024) found that 80% of cells recruited the PI(3,5)P2 sensor SnxA-GFP to a single location that they proposed was a membrane contact site, while in the absence of salt stress, SnxA-GFP was distributed evenly on the vacuolar membrane or in the cytosol. They also noted that the site of accumulation upon salt stress appeared to correlate with a site of vacuolar membrane invagination as vacuoles fragmented. In Figure 5C, we show that a duplicated Vph1NTNT-GFP becomes enriched at sites of invagination, further supporting these results. The association between PI(3,5)P2 synthesis and membrane invagination is not surprising, given that hyperactive alleles of *FAB1* result in extensive vacuolar fragmentation (Duex *et al*., 2006a; Lang *et al*., 2017), and reduced PI(3,5)P2 synthesis inhibits fragmentation and results in vacuole enlargement (Bonangelino *et al*., 2002; Efe *et al*., 2007).

Domains in the vacuolar membrane have been observed in response to numerous stresses (Leveille *et al*., 2022; Reinhard *et al*., 2023), and electron microscopy images revealed that under hyperosmotic stress (900 mM salt), domains enriched in PI(3,5)P2 were partially separated from V-ATPase-enriched domains (Takatori *et al*., 2016). More recently, vacuolar fission was shown to occur at nuclear-vacuolar junctions, contact sites between vacuoles and the perinuclear endoplasmic reticulum, which forms the nuclear membrane (Hanaoka *et al*., 2024). A requirement for a membrane-membrane contact site could account for the fixed location of Vph1NT recruitment in Figure 5A. Notably, both SnxA (Huda *et al*., 2024) and Vph1NTNT exhibit prolonged vacuolar membrane localization compared to Vph1NT-GFP. This may reflect the lower affinity of Vph1NT-GFP for PI(3,5)P2, which causes its dissociation from the membrane as the lipid diffuses away from its site of synthesis and/or lipid domains disappear with reduced stress. It is consistent with this model that Ivy1, a Fab1 inhibitor, appears to move laterally away from the site of Vph1NT-GFP recruitment, upon salt addition. Taken together, the data suggest that PI(3,5)P2 synthesis occurs at a focused site on the vacuolar membrane that has limited overlap with V-ATPase complexes. However, PI(3,5)P2 diffuses laterally in the membrane, allowing it to bind to its targets, including the V-ATPase.

The central role of Hog1 in the response to hyperosmotic stress has been examined extensively (Brewster *et al*., 1993; Posas *et al*., 2000; Hohmann., 2002; Westfall *et al*., 2004). However, Jin et al. highlighted the importance of PI(3,5)P2 synthesis at the yeast vacuole in the cellular survival of hyperosmotic stress, particularly salt stress (Jin *et al*., 2017). Specifically, a much larger proportion of *hog1Δ* than *fab1Δ* cells survives after the first four hours of salt stress, suggesting early synthesis of PI(3,5)P2 is critical for viability (Jin *et al*., 2017). We show that relocalization of Vph1NT-GFP is significantly strengthened and extended in a *hog1Δ* mutant (Fig. 6A), suggesting increased PI(3,5)P2 synthesis or delayed degradation. Hog1 is rapidly phosphorylated and activated under salt stress, and several Hog1 targets in the cytosol, including Gpd1 and other targets that promote glycerol production, are rapidly phosphorylated, even in the absence of a transcriptional response (Westfall *et al*., 2008). These responses may provide early osmoadaptation, and it is possible that the failure of Hog1 to phosphorylate these early cytoplasmic targets results in a stronger and more sustained signal for PI(3,5)P2 synthesis. Significantly, the results in Figure 6A argue against increased PI(3,5)P2 synthesis requiring the HOG pathway. The signaling pathways connecting salt stress to PI(3,5)P2 synthesis are still not completely clear, but the non-canonical cyclin-dependent kinase Pho85 is implicated in this process (Jin *et al*., 2016, 2017).

As described above, it is very likely that the V-ATPase activity is a major target of PI(3,5)P2 under salt stress, but it cannot replace the activities of Hog1. Consistent with this, multiple synthetic interactions between V-ATPase mutations and *hog1Δ* have been reported (Li *et al*., 2012; Banerjee *et al*., 2019). In Figure 6B, both *vph1Δ* and *hog1Δ* show growth defects during 20 h of salt stress, suggesting that even if PI(3,5)P2 levels are elevated in the *hog1Δ* mutant, Vph1- containing V-ATPases cannot sustain rapid long-term growth in this mutant.

Figure 7 portrays a model for early events at the vacuolar membrane when cells are exposed to high salt. In the absence of salt stress, V-ATPases are present as both assembled, active complexes at the vacuole responsible for the maintenance of vacuolar acidification and as disassembled, inactive V1 and V0 subcomplexes. In a *VPH1*/Vph1NT-GFP heterozygous diploid (Figure 2), Vph1NT-GFP is cytosolic. Exposure to high salt results in a rapid influx of Na^+^ into the cell along with an efflux of water. PI(3,5)P2 is synthesized at a focused area of the vacuole, which can be recognized by Vph1NT-GFP. Na^+^ is sequestered in the vacuole through the activity of exchangers, and the V-ATPase is assembled and activated through interactions with PI(3,5)P2 diffusing away from the site of synthesis, providing continued maintenance of vacuolar pH. As osmoadaptation proceeds, the Na^+^ is exported from the cell through newly synthesized channels, and balanced V-ATPase activity is restored.

**Figure 7:**
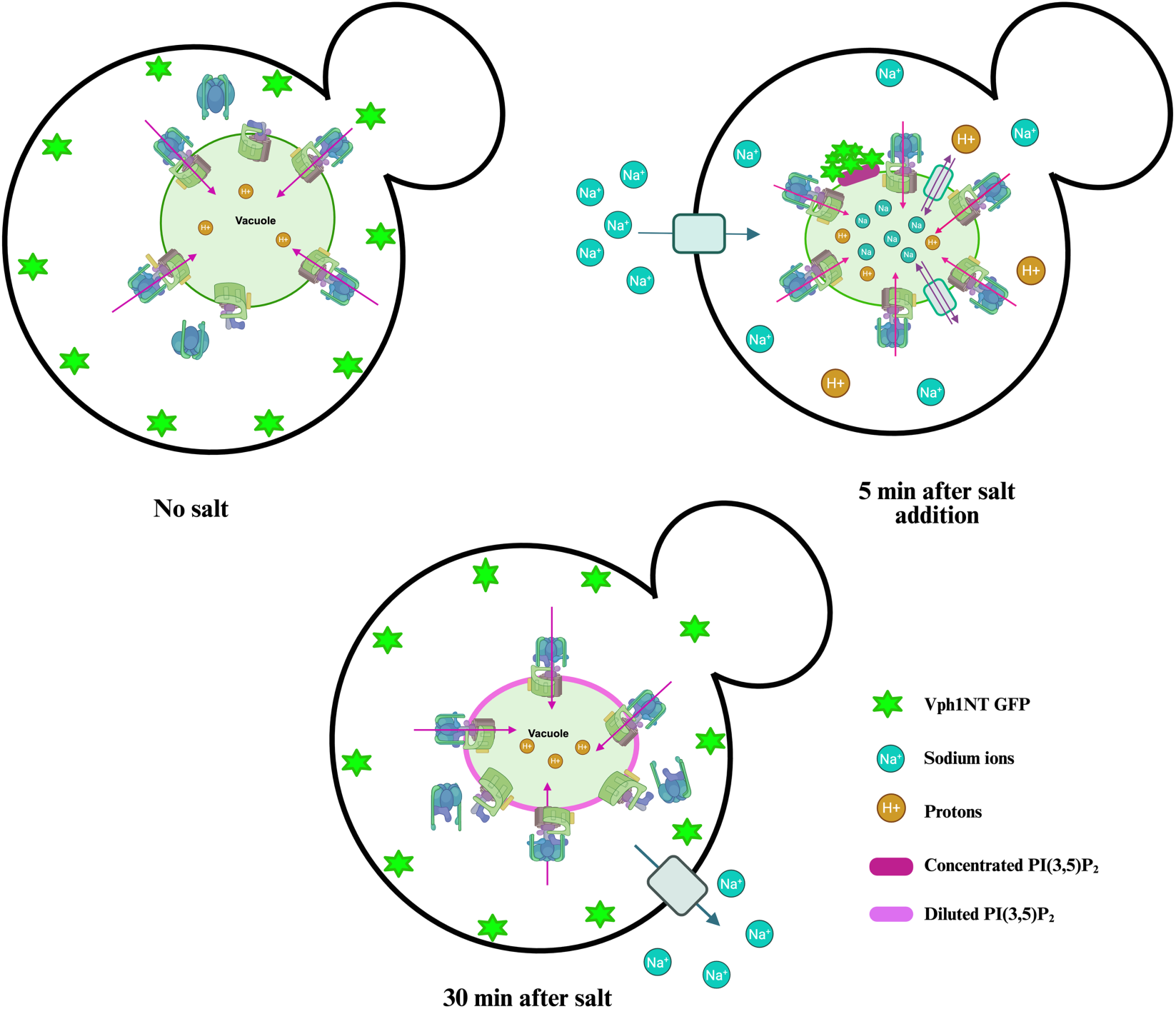
Model depicting early vacuolar responses of yeast cells to the addition of NaCl. Top left: Cells before salt addition. Vacuoles are acidified (brown circles in light green vacuole represent H^+^) by assembled V-ATPases, though some V-ATPases are typically disassembled. Vph1NT-GFP (Green stars) is cytosolic. Pink arrows indicate the proton transport from the cytosol into the vacuole by the V-ATPase. Top right: Cells about 4 min after salt addition. NaCl (light blue circles) enters the cytosol from the extracellular space and is sequestered in the vacuole by Na^+^/H^+^ exchangers. The V-ATPase is further assembled and activated to maintain acidification while also driving Na^+^/H^+^ exchange (Li et al, 2014). New synthesis of PI(3,5)P2 (magenta region of vacuole) is initiated at a region of the vacuole identified by Vph1NT-GFP clustering, before the lipid diffuses to the rest of the vacuolar membrane for V-ATPase activation. Bottom: Cells about 30 min after salt addition. Cytosolic levels of Na^+^ drop as newly synthesized Na^+^ exporters, upregulated by the Hog pathway, arrive at the plasma membrane. The V-ATPase returns to the resting state, and the vacuole remains acidic. Some Na^+^ may be stored in the vacuole. Vph1NT-GFP has returned to the cytosol.

Significantly, the major players investigated here are present in other eukaryotic cells. V-ATPases are very highly conserved, and we have established that a mammalian lysosomal a- subunit isoform also binds to PI(3,5)P2 (Mitra *et al*., 2023). The complex that synthesizes PI(3,5)P2 is also conserved (Rivero-Ríos and Weisman, 2022), and PI(3,5)P2 synthesis in response to salt stress has been reported (Jin *et al*., 2017). Given these common elements, it is certainly possible that PI(3,5)P2 is synthesized in focused domains of membranes in response to stress in other cell types.

## MATERIALS AND METHODS

### Materials

Oligonucleotides were purchased from Eurofins Genomics. PCR reactions were conducted using Q5 polymerase (New England Biolabs) and Accuris Taq polymerase (Accuris Life Science Reagents). Haploid and heterozygous diploid yeast *kanMX4*-knockout strains in the BY4741, BY4742, and BY4743 backgrounds were from knockout collections available from Horizon/Dharmacon. All media were purchased from Fisher Bioreagents. Other reagents were from Sigma unless otherwise indicated.

### Media

Synthetic complete (SC) medium was supplemented with 2% dextrose (Amberg *et al*., 2005). Where indicated, yeast extract-peptone-dextrose (YPD) medium was buffered to pH 5.0 with 50 mM potassium phosphate and 50 mM potassium succinate (Yamashiro *et al*., 1990). For the selection of the kanamycin-resistant (*kan^R^*) yeast strains, unbuffered YPD was supplemented with 200 µg/ml G418 (GoldBio). 100 µg/ml nourseothricin sulfate (GoldBio) was added to YPD for selection of *nat^R^* strains, and 200 µg/ml hygromycin B (Invitrogen) was added for *hyg^R^* strains.

For the selection of bacteria containing plasmids with an ampicillin-resistant marker, cells were cultured in Luria broth (pH 7.0) containing 125 µg/mL ampicillin (GoldBio).

### Construction of Yeast Strains

The Vph1NT-GFP-*kan^R^* construct consists of GFP and a *kan^R^* marker following the sequence for amino acid 406 of Vph1p that was constructed as described (Li et al., 2014). All yeast strains used in this study are listed in **Table 1**, and the plasmids used in this study are listed in **Table 2**. For genomic integration into other strains, the construct was amplified by PCR and introduced into the cells by transformation (Gietz and Schiestl, 2007). Transformants were selected on YEPD +G418 plates and confirmed by PCR from yeast genomic DNA (Masterpure yeast DNA Purification Kit). The 5Aa-Vph1-mCherry::hph1Mx6 (*hyg^R^*) cell line was created by using the pBS35 plasmid (a gift from Eric Muller) (Hailey *et al*., 2002) as a template, and amplified with oligonucleotides Vph1-mCherry HYG F and Vph1FL-mCherry HYG R. (See **Table 3** for oligonucleotide sequences) The resulting PCR product was transformed into SF838-5Aa cells, and transformants were selected on YEPD + hygromycin. Vph1NT-GFP::*nat^R^*was generated by switching the *kan^R^* marker using the p4339 plasmid as a template, as described by (Tong *et al*., 2001) using Nat R Forward and Nat R Reverse oligonucleotides. The PCR product was transformed into SF838-5Aα-Vph1NT-GFP-*kan^R^* cells. Transformants were selected by growth on YEPD + nourseothricin plates, and loss of G418 resistance was confirmed. To create a diploid strain heterozygous for Vph1NT-GFP-*kan^R^* and full-length Vph1 or Vph1-mCherry, haploid SF838-5Aα-Vph1NT-GFP-*kan^R^* was mated to either a wild-type SF838-5Aa strain or SF838-5Aa Vph1-mCherry-hphMx6, and zygotes were selected by micromanipulation after ∼7 hours of mating. To generate a *hog1Δ::kan^R^* strain with Vph1NT-GFP *nat^R^*, we made the diploid by mating BY4741 *hog1Δ::kan^R^* and BY4742-Vph1NT-GFP::*nat^R^* strains. Diploids were selected on plates containing both antibiotics and tetrad dissection was performed to obtain haploid spores. A haploid *hog1Δ*::*kan^R^* Vph1NT-GFP::*nat^R^* strain was then used for imaging. Construction of the Vph1NTNT-GFP strain, which contains a tandem duplication of amino acids 1-406 of Vph1 followed by a single GFP, was described previously (Li et al., 2014). Diploid *vac14Δ/vac14Δ* Vph1NT-GFP/*VPH1* strains were generated by mating BY4741*vac14Δ::kan^R^* Vph1NT-GFP::*nat^R^* with BY4742 *vac14Δ::kan^R^*, and sporulating the resulting diploid. Similarly, diploid strains containing *hog1Δ*::*kan^R^*and Vph1NT-GFP::*nat^R^* or *hog1Δ::kan^R^* and *VPH1* were generated by mating BY4741 *hog1Δ::kan^R^* Vph1NT-GFP::*nat^R^*or BY4741 wild-type cells with BY4742 *hog1Δ::kan^R^* and selecting zygotes under a dissecting microscope for further imaging. The 5Aa- Ivy1-mCherry:: *kan^R^* cell line was created by using the pBS34 plasmid (mCherry-kanMx4), a gift from Eric Muller (Hailey *et al*., 2002) as a template, and amplified with oligonucleotides IVY1- MCHERRY-FWD-F2 and IVY1-MCHERRY-REV-R1. The resulting PCR product was transformed into SF838-5Aa cells, and transformants were selected on YEPD +G418. To create a diploid strain, 5Aα-Vph1NT-GFP-*kan^R^* and 5Aa-Ivy-mCherry-*kan^R^*were mated, and diploids were selected for imaging.

**Table 1:**
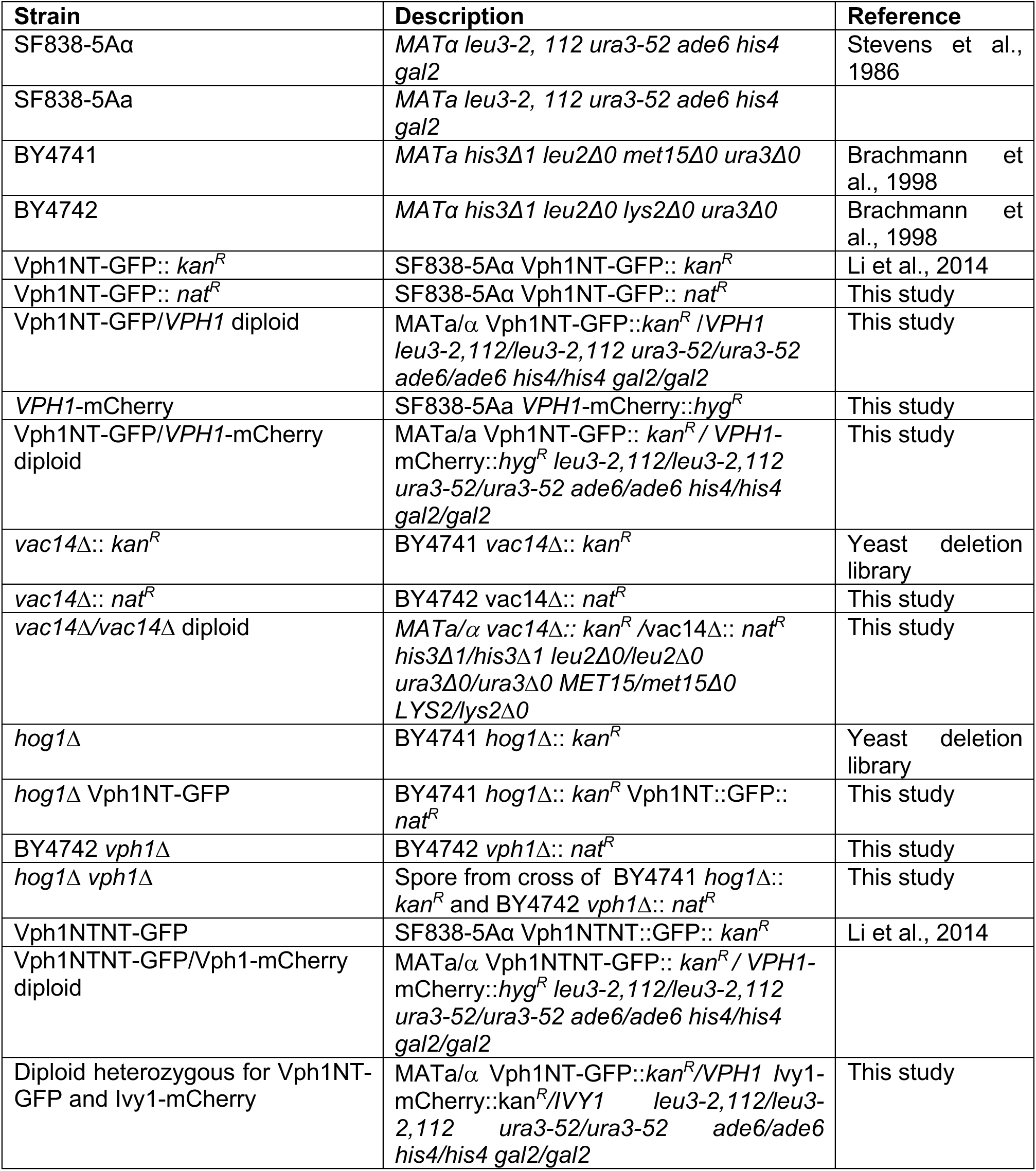
List of Yeast strains used in the study.

**Table 2:**
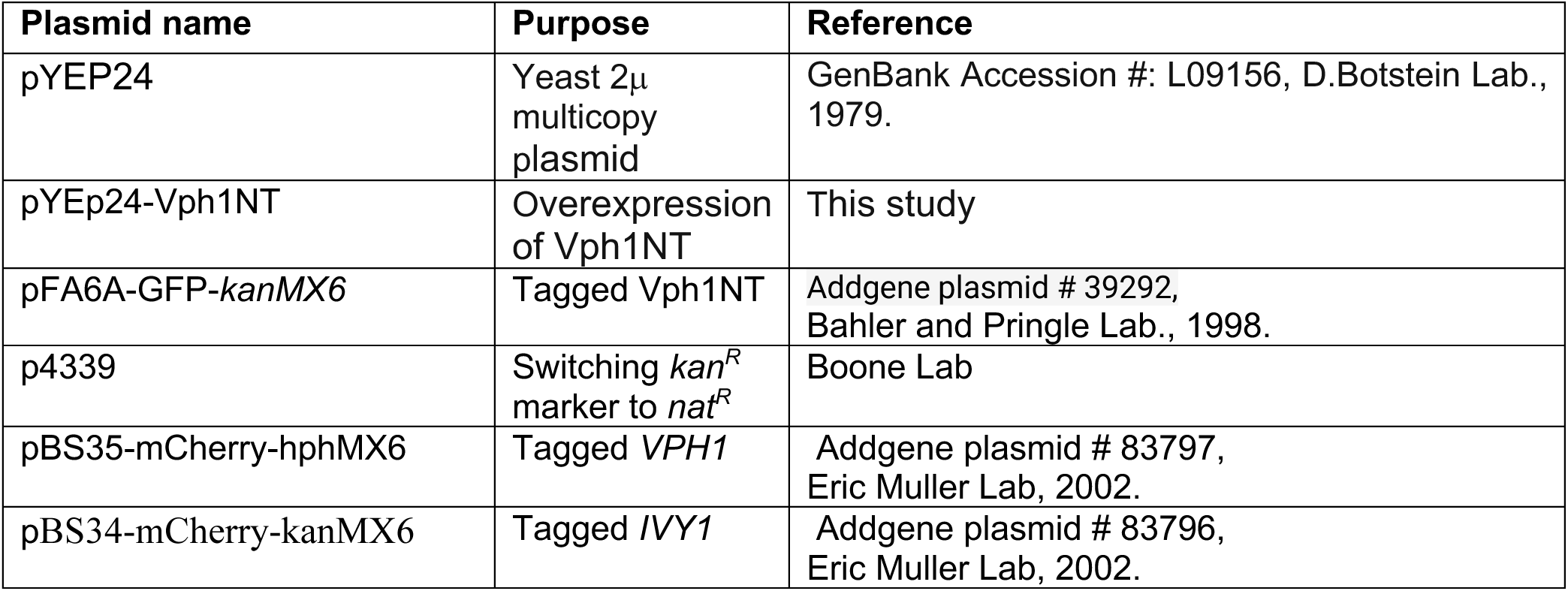
List of Plasmids used in the study.

**Table 3:**
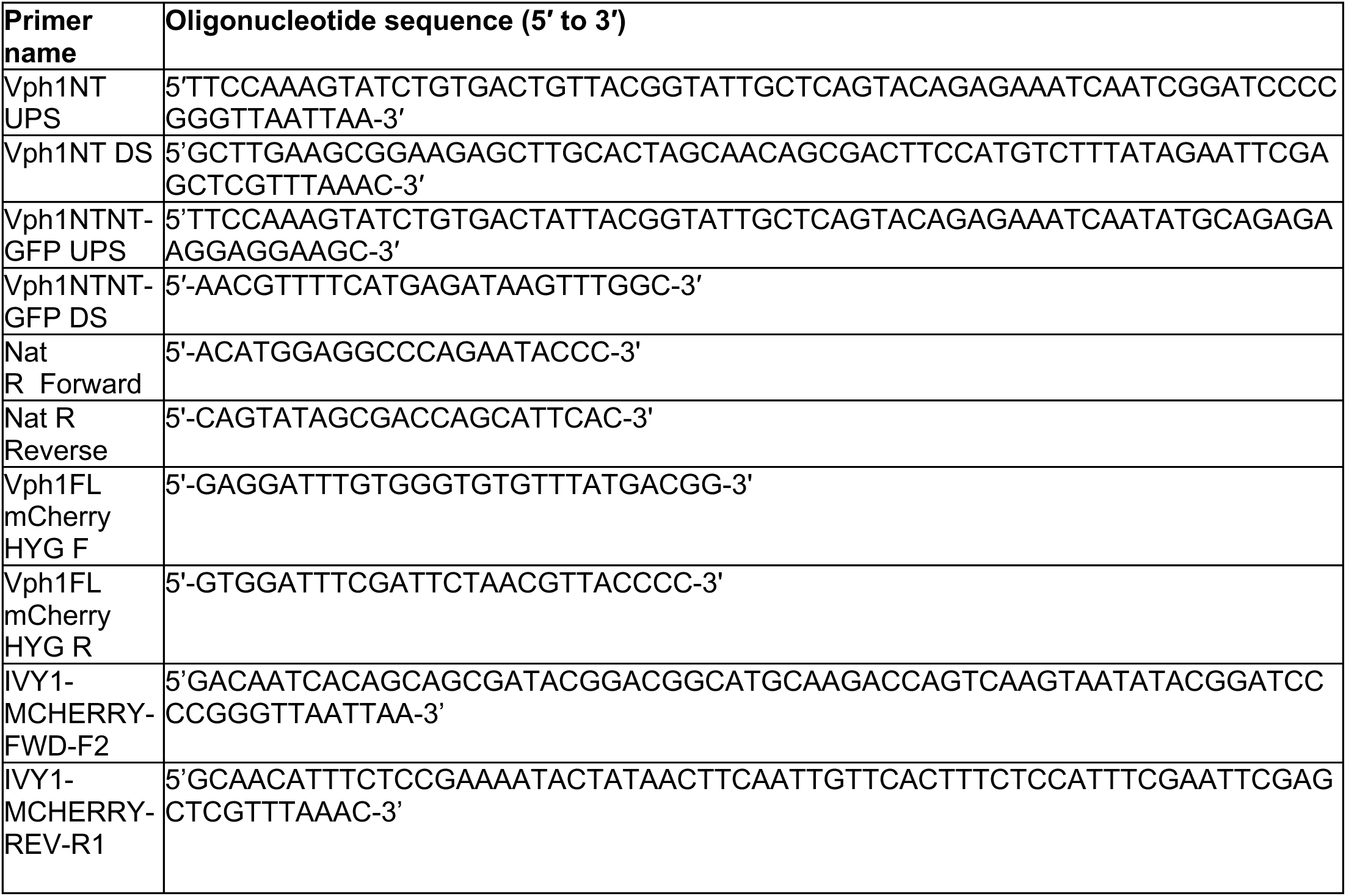
List of Oligonucleotides used in the study.

To obtain strains for comparison of growth rates under salt stress, BY4742 *vph1Δ:: nat^R^*, was mated with BY4741 *hog1Δ::kan^R^.* To generate a BY4742 *vph1Δ::nat^R^*, a heterozygous *vph1Δ::kan^R^* strain from the yeast heterozygous diploid library was sporulated, and a MATa spore containing *vph1Δ::kan^R^* was selected. The *kan^R^* marker was then switched to *nat^R^*as described above. The BY4741 *hog1Δ::kan^R^* mutant was obtained from the yeast haploid deletion library and crossed to a BY4742 *vph1Δ*::*nat^R^* strain. After sporulation of the diploid and tetrad dissection, four haploid spores with wild type, *hog1Δ, vph1Δ*, and *hog1Δvph1Δ* genotypes were selected for growth assays. All cells were grown overnight in SC medium, then diluted to an OD600 of 0.01 in SC containing either no salt or 500 mM NaCl, transferred to a 96-well plate, and placed in a SpectraMax i3X multi-mode multi-plate reader for growth at 30°C. Absorbance at 600 nm was recorded every 30 min over a total of 20 hrs. Three biological replicates derived from separate colonies were assessed for each strain.

### Overexpression of Vph1NT

We have used the YEp24 vector (Table 2), a yeast 2μ shuttle vector, for Vph1NT overexpression in *Saccharomyces cerevisiae* (Botstein *et al*., 1979). The YEp24 plasmid containing Vph1NT was introduced into haploid (SF838-5Aα) or diploid cells heterozygous for Vph1NT-GFP*::kan^R^* and Vph1FL-mCherry::*hyg^R^*. Transformants were selected on SC medium lacking uracil, then grown overnight to log phase. After pelleting and washing with fresh media, they were imaged on a Zeiss Axio Imager Z1 widefield microscope equipped with a Hamamatsu ORCA-ER CCD camera and a 100x oil (NA 1.4) objective and DIC optics. Image acquisition was done using AxioVision 4.8 software (Carl Zeiss, Peabody, MA). Vacuole diameters were measured for 70-90 cells per biological replicate, using the line scan tool in FIJI. The data points for three biological replicates were plotted using GraphPad Prism. Statistical analysis was performed using an unpaired t-test, and data are presented as mean ± SEM.

### Microfluidics setup and fluorescence Imaging

Microfluidic experiments were conducted using the ibidi µ-Slide VI 0.1 uncoated six-channel chambers (1.7 μL channel volume, 60 μL reservoir volume). Channels were precoated with concanavalin A (final concentration 1 mg/ml) to facilitate yeast cell adhesion and washed with water after 30 minutes. The indicated strains expressing GFP-tagged Vph1NT were grown overnight to log phase in SC medium, pelleted, and resuspended in fresh medium for imaging. Cells were introduced into the chamber via a motorized pump (Harvard Apparatus) operating at 10 μl/min, allowing a controlled flow of cells, media, and NaCl. Time-lapse imaging began once the cells had settled. Images from a single central focal plane were captured using a Nikon Ti2-E SoRa spinning disk confocal microscope with a 60x 1.4NA oil immersion objective. GFP and mCherry fluorescence were visualized using 100 mW 488 nm and 100 mW 561 nm lasers, respectively. All the samples were imaged at GFP at 300 ms and 45% laser intensity, whereas mCherry was imaged at 100 ms and 25% laser intensity. The Nikon Denoise software module was applied to each image unless indicated otherwise.

Unless otherwise specified, experiments were conducted as 15-minute time-lapse videos with images captured every 30 seconds to give 31 time points/experiment. For the first 3 minutes, the cells were imaged without salt to set a baseline, salt was then added, and imaging continued for the remainder of the loop. For diploid strains expressing both GFP and mCherry, imaging conditions were identical to those for haploid cells, with images obtained at both excitation wavelengths every 30 seconds. Reversibility experiments began with no salt for 3 minutes, then salt was added and incubation continued for 12.5 minutes. At minute 13, medium without salt was added and the incubation was continued for 7 minutes, followed by readdition of salt at 20 minutes for the remaining time. For haploid and diploid *hog1Δ* mutants, both wild-type and mutant strains were imaged for a total of 30-40 minutes, with salt addition after 3 minutes. Images were obtained every 30 seconds.

### Image Processing and Quantification

For each biological replicate, fluorescence intensity across time points 1 to 31 was quantified by first selecting 8-10 cells present at all time points, then drawing a line across the same position at each time point, and quantifying fluorescence using the linescan function in FIJI. The resulting fluorescence intensity plot revealed peaks at the vacuolar membrane, which were quantified as the peak intensity (maximum fluorescence) to assess Vph1NT-GFP localization at different time points. Maximum fluorescence intensity was normalized to the intensity at pre-salt time points, averaged, and plotted for each biological replicate. At least three biological replicates, corresponding to cells cultured from distinct colonies, were analyzed for each condition. Statistical analysis was conducted with GraphPad Prism 9 using two-way ANOVA with repeated measurements.

The percentage of cells with Vph1NT GFP relocalization was scored manually at each of the 31 time points across three biological replicates and different salt concentrations. 250-300 cells were counted per biological replicate. Statistical analysis was conducted with GraphPad Prism 9 using two-way ANOVA with repeated measures. For comparing differences between wild-type and mutant cell lines, a multiple paired t-test was used.

A minimum of three biological replicates were conducted for each condition. Data are presented as mean ± SEM. The percent recruitment and normalized maximum fluorescence intensity of Vph1NT-GFP recruitment were plotted for all time points from a representative sample. Bar graphs showing results from three biological replicates at specific time points are also included in Figures 1-3. Two-way ANOVA with repeated measurements was used to compare values across different salt concentrations; comparisons with a p-value < 0.05 were considered significant.

## ACKNOWLEDGMENTS

This work was supported by the National Institute of Health Grant NIH R35 GM145256 to P.M.K. We thank Jessica Henty-Ridilla for helping with microscopy, Wenyi Feng for helping with the statistical analysis, and Maureen Tarsio for helping with media and experiments. We also thank Harsimranjit Sekhon for writing a programming code for the image analysis.

Illustrations were prepared using Biorender.com.

## Conflict of interest

The authors declare that they have no conflicts of interest with the contents of this article.

## Data availability

Any data and materials will be shared upon request by contacting Patricia Kane. Source data supporting this study’s findings will be available at https://upstate.figshare.com (doi: 10.58120/upstate.29856158)

## Author contributions

K.S. performed experiments, analyzed data, prepared figures, and wrote the first draft of the manuscript; P.M.K. obtained funding for the project, analyzed data, prepared figures, and contributed to the writing of the manuscript.

## List of Abbreviations used in the study

Vph1NT: N-terminal cytosolic domain (aa 1-406) of Vph1p
Vph1CT: C-terminal membrane domain of Vph1p
PI: Phosphatidylinositol lipids
PI(3,5)P_2_: Phosphatidylinositol 3,5-bisphosphate
HOG pathway: High Osmolarity Glycerol pathway
DIC: Differential interference contrast
GFP: Green Fluorescent Protein
V-ATPase: Vacuolar proton-translocating ATPases

## Notes

### Competing Interest Statement

The authors have declared no competing interest.

