## Supplemental Figure 1 for "Early responses to hyperosmotic stress at the yeast vacuole"

**Supplementary Figure 1: Vph1NT GFP integrated and expressed from the *VPH1* locus does not affect vacuolar diameter.**

Left: DIC images of haploid *vph1*Δ and Vph1NT GFP (integrated and expressed from the *VPH1* locus as in Figure 1). Scale bar = 4 μm. Right: The largest vacuole diameter in haploid *vph1*Δ and Vph1NT GFP cells was determined in 50-75 cells per biological replicate, as in Figure 3. Dots represent all vacuolar diameters quantitated across the three replicates, along with the mean ± SEM. ns = not significant as calculated by unpaired Student's t-test.

### Supp. Figure 1

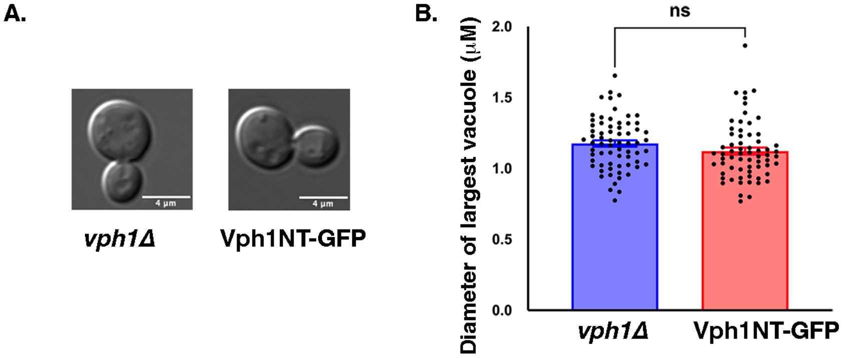
